# Re-expressing coefficients from regression models for inclusion in a meta-analysis

**DOI:** 10.1101/2021.11.02.466931

**Authors:** Matthew W. Linakis, Cynthia Van Landingham, Alessandro Gasparini, Matthew P. Longnecker

## Abstract

Meta-analysis poses a challenge when original study results have been expressed in a non-uniform manner, such as when regression results from some original studies were based on a log-transformed key independent variable while in others no transformation was used. Methods of re-expressing regression coefficients to generate comparable results across studies regardless of data transformation have recently been developed. We examined the relative bias of three re-expression methods using simulations and 15 real data examples where the independent variable had a skewed distribution.

Regression coefficients from models with log-transformed independent variables were re-expressed as though they were based on an untransformed variable. We compared the re-expressed coefficients to those from a model fit to the untransformed variable. In the simulated and real data, all three reexpression methods usually gave biased results, and the skewness of the independent variable predicted the amount of bias. Meta-analyses stratified by whether the key independent variable was log-transformed and synthesis of the stratified results without meta-analysis appear to be preferable to use of the re-expression methods examined.

## 1. Introduction

The results of a group of studies deemed comparable can be synthesized quantitatively using meta-analysis. To base the meta-analysis on all available data, “Results extracted from study reports may need to be converted to a consistent, or usable, format for analysis” (J. P. Higgins et al., 2019). Methods of converting data presented by authors into a format suitable for meta-analysis have been well developed for intervention studies, but for studies of exposure effects the available tools are somewhat limited.

A challenge that frequently arises in the meta-analysis of non-randomized studies of exposure effects is that the authors of the studies being synthesized have made different choices about how to represent a continuous-scale exposure in their analysis. For example, the exposure might be left in its original untransformed units, log-transformed, or categorized.

Various approaches to the problem of inconsistently-expressed effect estimates have been recommended or used in practice (Campbell et al., 2020; McCabe et al., 2021; Negri et al., 2017; Ye et al., 2019). Obtaining the raw data or asking authors to re-analyze their data are the ideal solutions, though not always practical. When these options are not feasible, the results from studies using the lessfrequent approach have been be excluded from the meta-analysis (McCabe et al., 2021), or preferably the results of studies that used transformed and original units are analyzed separately (Negri et al., 2017; Ye et al., 2019), and then synthesized without meta-analysis (SWiM) (Campbell et al., 2020). Some authors, however, have recently used re-expression methods to address the problem (Dzierlenga et al., 2020; Steenland et al., 2018). The validity of these re-expression methods, however, have not been evaluated in detail. Here we consider methods of re-expressing regression coefficients from linear models fit to a log-transformed exposure variable as the coefficient that would have been obtained had the authors left the exposure in its original units. We refer to this process as re-expression of *β* to an untransformed basis.

An algebraic method of re-expressing regression coefficients was recently described and evaluated using one simulated data set and one set of parameters (Rodríguez-Barranco et al., 2017). Rodriguez-Barranco et al. found that in the setting of a log-transformed lognormally distributed independent variable, when the beta coefficient from a model fit to the transformed data was reexpressed to what they would have gotten had the model been fit to the untransformed data, the reexpressed coefficient was half the size of the true (fitted) coefficient. They recommended caution in applying their method when the distribution of the independent variable was markedly asymmetric. More recently, authors have developed computational methods of re-expressing coefficients from models fit to a log-transformed independent variable to approximate what would have been obtained if the model had been fit with the original unit continuous independent variable (Dzierlenga et al., 2020; Steenland et al., 2018). The basic principle is to minimize the difference between the *y* predicted from *y* = *β*·log(*x*) and the *y* predicted from a *y* = *β*·*x* (over the same range of *x*) by varying *β* in the second equation. Figure 1 may aid visualization of the task, where *y* from *y* = *β*·log(*x*) is shown with a light blue dotted line, and the difference in *y* from a straight line is minimized over a range of *x*. When Steenland et al. originally described the procedure it was for a fixed range of *x*, applicable to a specific exposure, and the validity of their method was not evaluated. Dzierlenga et al. (2020) used the same basic principle as Steenland et al. with a modification of the method to be more flexible with respect to the range of the exposure variable and found that it performed well when evaluated using data from five studies of one exposure. In addition to the above re-expression methods, we developed a third (“Alternative”) estimator that is algebraic but different than that of Rodriguez-Barranco et al., and introduce it below, in the methods section.

**Figure 1:**
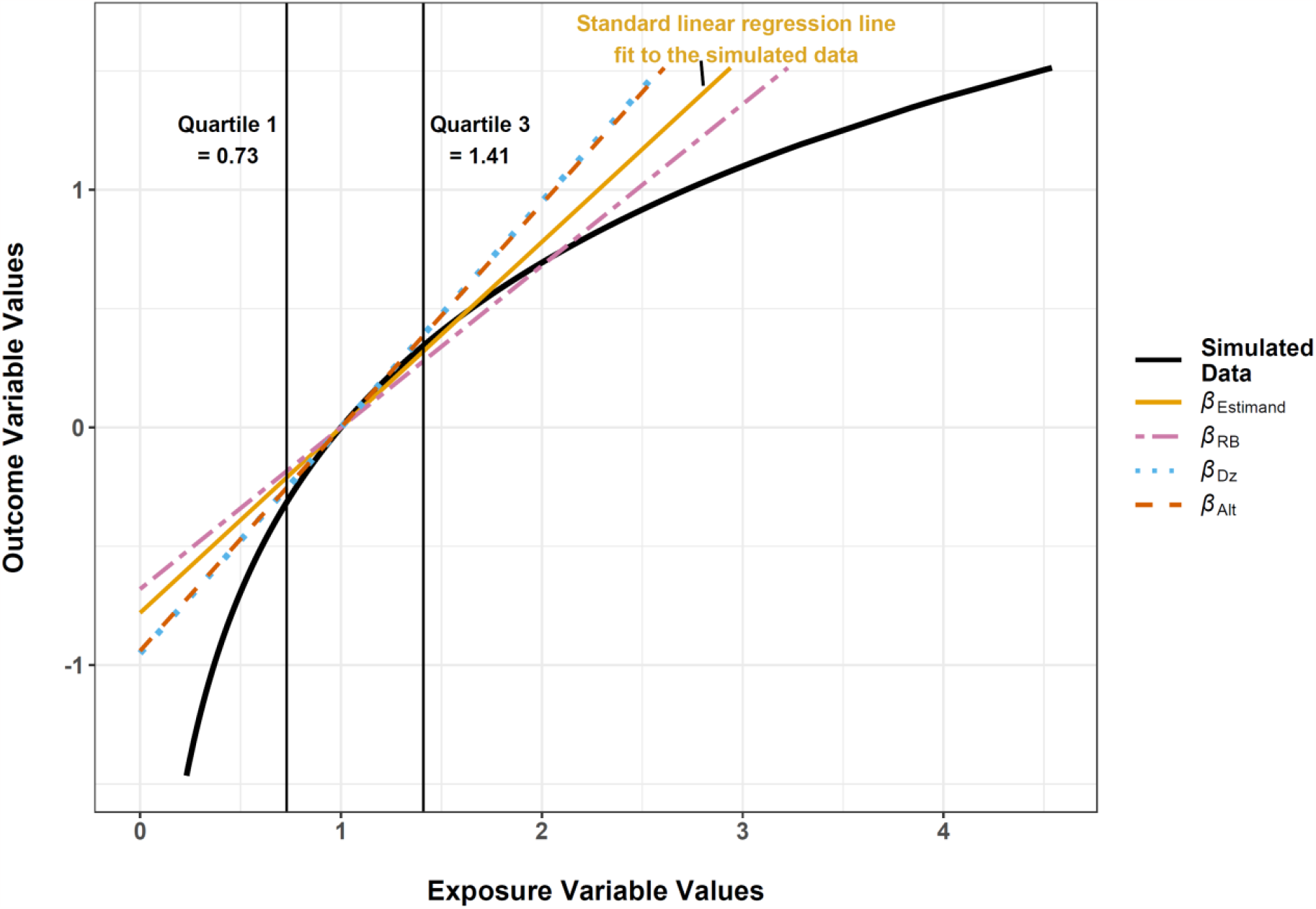
Plot of simulated values of *y* as a function of *x* from *y* = ln(*x*)(curve), along with slopes obtained by four methods (diagonal lines). The gold solid line represents aslope (*β*_Estimand_) fitted with the model y = α + βx. The three dashed lines are estimates of *β*_Estimand_ obtained by the re-expression methods described in the text. Vertical lines indicate the first and third quartiles of the x-values. The intercepts of the diagonal lines have been adjusted to emphasize the similarity of the slopes in the interquartile range.

The goal of the present project was to evaluate the validity of re-expression of regression coefficients to an untransformed basis for three methods using a wide variety of simulated and real data examples.

## 2 Methods

In this section, we present the simulation study that was used to evaluate the three estimators, and then describe the real datasets that were used to further evaluate the estimators. Our description of the simulation study follows the “ADEMP” format recommended by Morris et al. (2019), where ADEMP stands for Aims, Data generating mechanism, Estimand (target of analysis), Methods, and Performance measures (Morris et al., 2019). The methods subsection of the ADEMP gives a detailed specification of the estimators and is thus relatively long.

### 2.1 Description of the simulation in ADEMP format

Aim: To examine bias in the estimated regression coefficients (regression coefficient that would have been obtained had the original analysts not log-transformed exposure before fitting a regression model) calculated by three methods.

Data generating mechanism (DGM): An independent random variable *x* with a log normal distribution used to define the dependent variable *y* = *β*_DGM_·log_*b*_(*x*) + *e*. The model parameters, possible values, and rationale for the chosen values are shown in Table 1. A *β*_DGM_ = 0 was not studied in the simulations because it caused instability in the relative bias performance measure. A factorial simulation design was used with the parameter values indicated in Table 1. Specifically, every possible combination of parameter values was used, for a total of 960 simulation scenarios (each with n_sim_ = 2000).

**Table 1.** Values of parameters used in the simulation and rationale for their choice. Number of simulations = 2000.

| Parameter | Possible values | Rationale for the choice |
| --- | --- | --- |
| $n_{\text{obs}}$ | 162, 8474 | Quantiles (10<sup>th</sup>, 90<sup>th</sup>) of the distribution of sample sizes for the 15 real data examples. |
| $e$ | Selected from a normal distribution with mean = 0 and SD = $SD_e^*$ | Standard deviation selected to result in an $R^2 \sim 0.2$, to make realistic models |
| $\beta_{\text{DGM}}$ | -15, 0.5, 1, 10, 30 | Broad range of effect sizes encompassing those in the real data examples. |
| Logbase | 2, e, 10 | Log bases used in the 15 real data examples. |
| $\sigma$ | 0.25, 0.45, 0.65, 0.85 | Selected values cover the approximate range of $\sigma$ values in the 15 real data examples. |
| median<br>($\mu = \log(\text{median})$) | 0.25, 0.5, 1, 2, 4, 8, 16, 32 | Selected values cover the range of median exposure in the 15 real data examples. |

\* $SD_e$ was calculated using the following method: $SD_e = |\beta_{\text{DGM}} * \log(\text{median} + 2) * \text{Mult}_{\text{SD}}|$ where $\beta_{\text{DGM}}$ and median are from the table above and $\text{Mult}_{\text{SD}}$ is a multiplier to prevent extremely large variability. For each median, the $\text{Mult}_{\text{SD}}$ was as follows: median 0.25, $\text{Mult}_{\text{SD}}$ 1; 0.5, 0.886; 1, 0.771; 2, 0.657; 4, 0.543; 8, 0.429; 16, 0.315; 32, 0.120). $\text{Mult}_{\text{SD}}$ was selected based on values that will introduce enough error to result in $r^2$ values around 0.2 (similar to many epidemiological studies) for estimation of $\beta_{\text{Estimand}}$ .

Estimand: The *β* coefficient from fitting *y* = *β*_Estimand_·*x* + *e* with an ordinary least squares (OLS) model.

Methods: The three estimators evaluated were 1) as described by Rodriguez-Barranco et al. (2017), 2) as described by Dzierlenga et al. (2020), and 3) an approach we introduce below and call the Alternative estimator. We refer to these as *β*_RB_, *β*_Dz_, and *β*_Alt_, respectively.

An algebraic method for re-expression of *β* to an untransformed exposure was first presented by Rodriguez-Barranco et al. (2017). Equation 1 below shows their formula (See Model B in Table 1 of their publication):

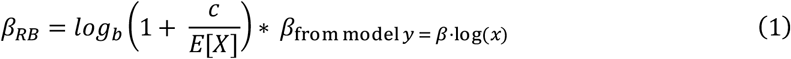

In equation 1, *β*_RB_ is the re-expressed beta coefficient using the Rodriguez-Barranco method, *b* is the log base used to transform *x, c* is the absolute change in exposure *x* (*c* = 1 unit of exposure in the present study), E[*X*] is the mean of the exposure, and *β*_*…*_ is the regression coefficient from the model using the log-transformed exposure. The same formula was applied to the confidence limits of *β* from the log(*x*) model.

A computational method for re-expression of *β* to an untransformed exposure was developed by Steenland et al. (2018), who described their method as

… iteratively minimizing the squared deviation of a new linear curve from the original logarithmic one, over a scale of 0 to 10 ng/ml PFOA [perfluorooctanoic acid], typical of studies in the general population. We also minimized squared deviation of a linear upper and lower confidence limit from the original logarithmic confidence interval curves. For any given study, the iteration was conducted by minimizing the sum of squares of the difference between the candidate linear curve and the logarithmic curve reported in the literature, across 10 points, at 1, 2…. through 10 ng/ml. Iteration began with an educated guess for a candidate linear curve that would approximate the logarithmic curves and proceeded by varying the candidate linear curve until the sum of squares of the differences were minimized.

Dzierlenga et al. (2020) used this same principle to calculate *β*_Dz_, though it modified it so that the method was more flexible with respect to the range of the exposure variable. The modification used an algorithmic optimization over 6 points from the 25th to the 75^th^ percentiles (25^th^, 35^th^…75^th^) of the estimated exposure distribution.

The Alternative method of algebraic re-expression that we developed for this report was based on the principle of calculating, on the untransformed scale of exposure, the increment that represented a doubling, a 2.718-fold increase, or a 10-fold increase (i.e., one log unit, with a base of 2, e, or 10). This was done by subtracting or adding 0.5 units on the log scale to the log(median exposure), backtransforming the results, and taking the difference (see Equation 2).

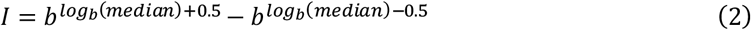

In equation 2, *I* is the increment used to re-express *β* from log to linear and *b* is the logarithm base. Then

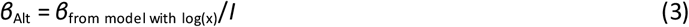

The same formula was applied to the confidence limits of *β* from the log(*x*) model. R scripts/functions and data files for applying each of these three re-expression methods are available in the supplemental materials (Supplemental_Code.zip).

Performance measures: We focused on relative bias, coverage probability, and the Monte Carlo standard error of the relative bias for each estimator. An example of the formula for the mean relative bias for a given scenario is:

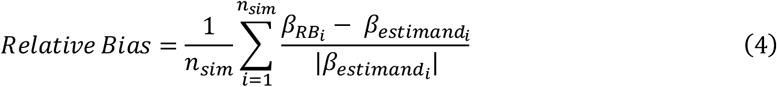

Where, e.g., 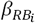refers to the beta coefficient obtained from the Rodriguez-Barranco et al. estimator for the *i*^th^ repetition, 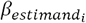refers to the beta coefficient obtained from the ordinary least squares estimator on the untransformed, simulated data, and *n*_sim_ is the number of simulations conducted.Absolute value of the β_estimand_ is used as the denominator in order to generate the correct sign for the absolute bias when both β values are negative. β_estimand_ is used rather than β_DGM_ to calculate the relative bias in equation 4 so that the results reflect the performance of the estimator(s) in specific datasets. An example formula for the Monte Carlo standard error of the relative bias for a given scenario is:

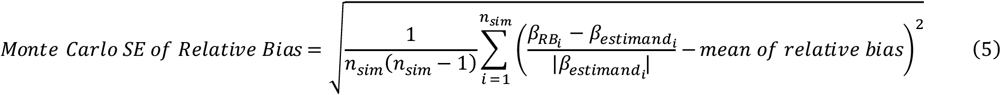

### 2.2 Evaluation of the determinants of relative bias using the simulated data

After running the simulations using the parameter values shown in Table 1, for each estimator we fit ordinary least squares models of the relative bias as a function of median, *σ, b* (log base), *n*_*obs*_, and *β*_*DGM*_, and interaction terms between sigma and these variables. A both-directions stepwise approach was taken where the multiple of the number of degrees of freedom used for the penalty (k) was set to a value ∼3.84 (p < 0.05 in Chi-square test) and the optimal model was selected by minimization of the AIC value. Each observation in the dataset analyzed was the average result from 2000 simulations. Use of the average rather than the data for all 1,920,000 (960·2000) observations resulted in essentially the same models and produced more interpretable plots.

### 2.3 Evaluation of the validity of the three estimators using the real data

To further evaluate the validity of re-expression methods and guide our simulations, we sought examples for various types of outcomes (dichotomous, log-continuous, untransformed continuous) and a variety of environmental agents with exposure measured using a biomarker. Environmental exposures measured with a biomarker frequently have skewed distributions with a long tail to the right. We first identified a series of published analyses based on data that were publicly available. Second, we identified a similar series of published analyses that did not have raw data available but that presented regression results obtained with and without log transformation of the exposure.

For the example data that involved our re-analysis of published results, we chose results that could efficiently be replicated to a reasonable degree of accuracy using the originally described methods. When the authors presented results for more than one outcome or more than one exposure in a report, in generalwe arbitrarily chose one result that was statistically significant for inclusion in our evaluation of validity; the exception was data from Xu et al. (2020), for which we included two results. Xu et al. (2020) showed results for two different outcomes, one continuous, and one dichotomous, that were examined in relation to the same exposure; the regression coefficients were statistically significant for both.

For each real data example, we calculated the relative bias for each of the three estimators (compared to the coefficient from the untransformed exposure), and then for the 15 examples calculated the median, quartiles, and range of relative bias values for each estimator.

In the two example datasets where the relative bias in the three estimators was largest, we explored whether the exclusion of influential observations affected the accuracy of the re-expression using *β*_Dz_. In two additional examples datasets where the relative bias was typical of other studies, we also examined the effect of excluding influential points on the validity of the re-expression with *β*_Dz_. Influential observations were identified with a difference in beta analysis (change in beta with each observation excluded one at a time) performed on the regression using untransformed exposure. A t-test-like statistic was used to identify the 5% of points that were unusually influential (|DFBETAS| > 2/√n) (Belsley et al., 1980).

### 2.4 Adjustment for bias in the estimators

The regression equations we developed to evaluate the determinants of relative bias in the simulated data (Section 2.2) were used to predict the relative bias in each estimator based on σ and other parameters, as needed. The predicted relative bias was used to estimate what the value of the estimator would have been were it not biased, e.g., *β*_Alt,adjusted_ = *β*_Alt_/(1 + predicted relative bias of *β*_Alt_). We applied this to the real datasets, to see if the adjustment resulted in an improved estimator.

## 3. Results

### 3.1 Simulations

A simplified example simulation with data generated by *y* = *β*_DGM_·log_e_(*x*) and parameters *β*_DGM_ = 1, median = 1, *σ* = 0.5, *SD*_e_ = 0 is depicted in Figure 1. In this scenario *β*_RB_ slightly undershot the slope estimated from the fitted regression line, whereas the *β*_Dz_ and *β*_Alt_ estimators overshot the fitted slope, by a slightly greater magnitude.

The relative bias of *β*_RB_ was a function of *σ* and the median exposure level (Figure 2A). When *x* was significantly skewed (e.g., *σ* = 0.65) and the median was 1, the relative bias was close to zero, but with other combinations of *σ* and median the range of bias was substantial. The coefficients for the model of relative bias in *β*_RB_ are shown in Table S1.

**Figure 2:**
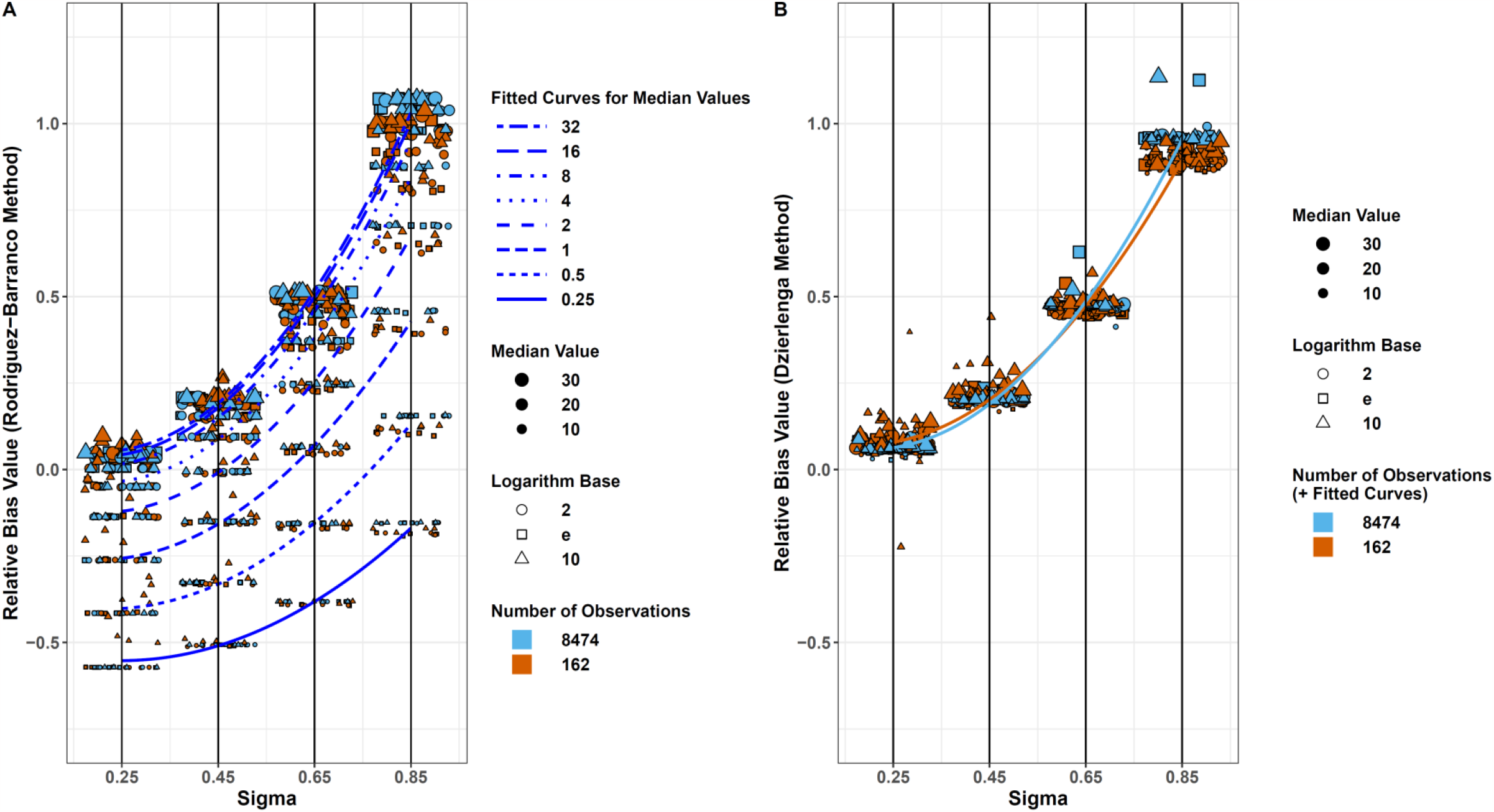
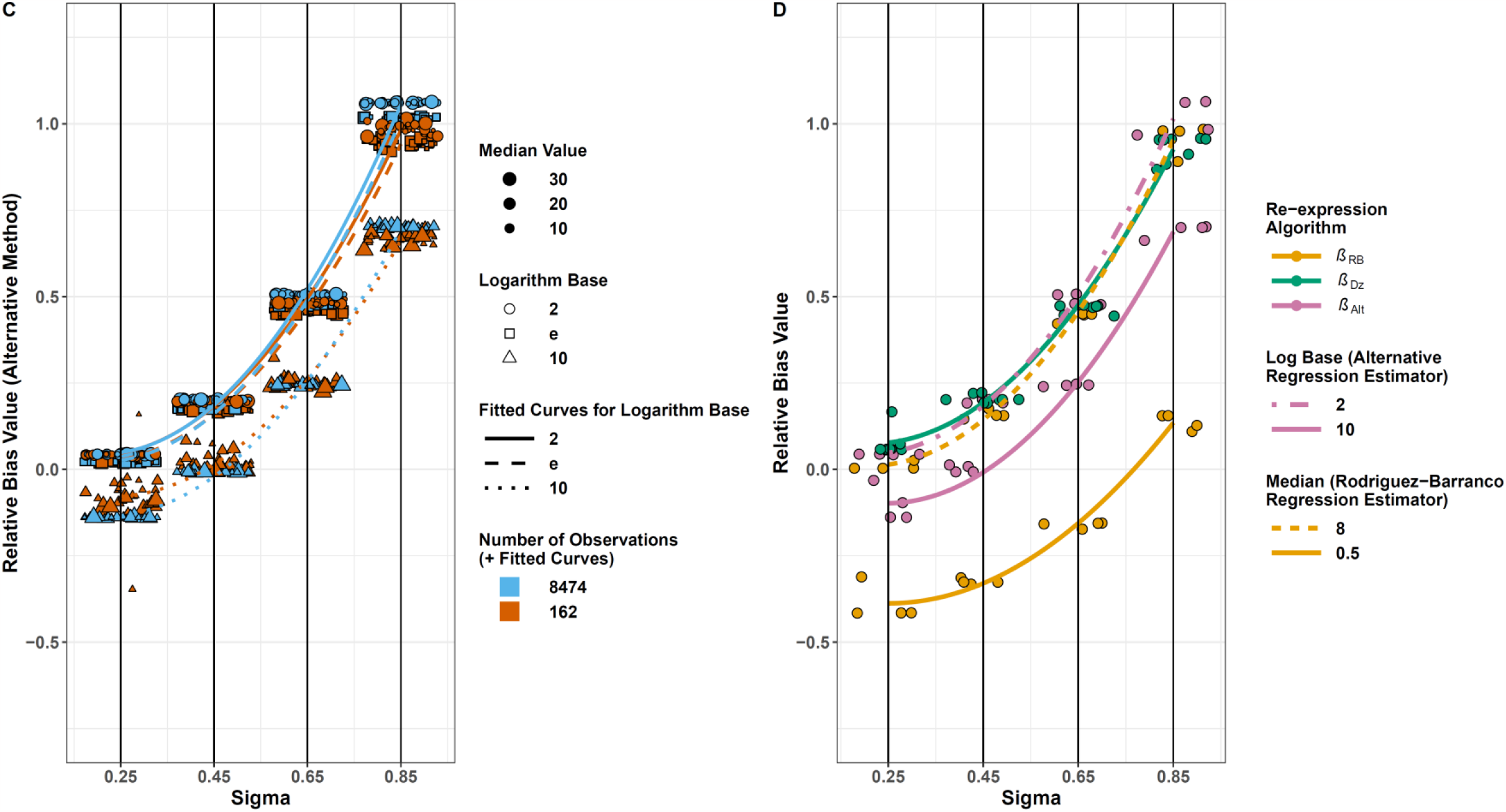
Plots of relative bias as a function of skewness (*σ*) in the exposure *x*, by type of estimator. Individual points represent the average result (*n*_sim_ = 2000) for each simulation scenario. A total of 890 of the possible 1,920,000 observations (960 scenarios x 2000 simulations) were not used in the calculation of the average results because *β*_estimand_ was < 0.0001 (essentially zero). Lines represent quadraticfits to the data for a specified prediction equation and set of values of independent variables (see text). Note that data have been artificially spread along the x-axis for visualization purposes, all actual x-values are the closest black vertical line (0.25, 0.45, 0.65, or 0.85). Figures A-C show points for 768 simulation scenarios (β_DGM_ > 0, see Figure S1 for plots including β_DGM_ < 0); Figure D shows points for a subset of scenarios (n = 32) chosen because they demonstrate differences among the estimator properties. A) Rodriguez-Barranco estimator, B) Dzierlenga estimator, and C) Alternative estimator. D) Shows all 3 estimators in the same plot with a subset of the data of the simulation data where *β*_DGM_ = 0.5, log base = 2 or 10, and median value = 0.5 or 8.

The relative bias of *β*_Dz_ was primarily a function of *σ*; the interaction of *σ* and *n*_obs_ was statistically significant but not substantially important (Figure 2B, Table S1). When *x* was significantly skewed (e.g., *σ* = 0.65), the relative bias was approximately 0.5.

The relative bias of *β*_Alt_ was primarily a function of *σ* and log base (Figure 2C, Table S1). When log base was 10, at a given value of *σ* the relative bias was lower than when log base was 2 or e. When log base was 2 or e, *β*_Alt_ performed similarly to *β*_Dz_.

As noted earlier, the models of relative bias for each estimator were fit to datasets with an n of 960, where each of the 960 observations was the average of 2000 simulations for each scenario (parameter set). When the same models were fit to all of the original data points (960·2000) the model fit statistics were essentially the same (Table S2).

Figure 2D shows the relative bias after restricting the parameter set for the simulation to best display the key properties of each estimator: all depend on *σ, β*_RB_ additionally depends on the median, and *β*_Alt_ additionally depends on the log base. The overall interpretation based on the figure was that in the simulations in general, with *σ* > 0.45 the estimators were substantially biased except for specific circumstances where *β*_RB_ did well. Another way to summarize the overall findings was by the performance measures presented in Table 2, based on the results for all simulation scenarios with |β_Estimand_| > 0. The median Monte Carlo standard error (MCSE) across all three estimators was 0.002. Thirty percent of the 2880 simulations (960 scenarios ·3 estimators) had an MCSE > 0.005. More than 95% of all 2880 simulations had an MCSE that was ≤ 0.02 (relatively small compared with the average relative bias). Among the < 5% with an MCSE > 0.02, the nobs was 162 and the log base was 10 in all instances. The maximum MCSEs were: β_RB_, 0.196; β_Dz_, 0.320; and β_Alt_, 0.262. The coverage probabilities were substantially below 95%, reflecting how infrequently the estimators performed well. Compared with *β*_RB_, the other two estimators, on average, had larger positive bias, but with higher coverage probabilities.

**Table 2:**
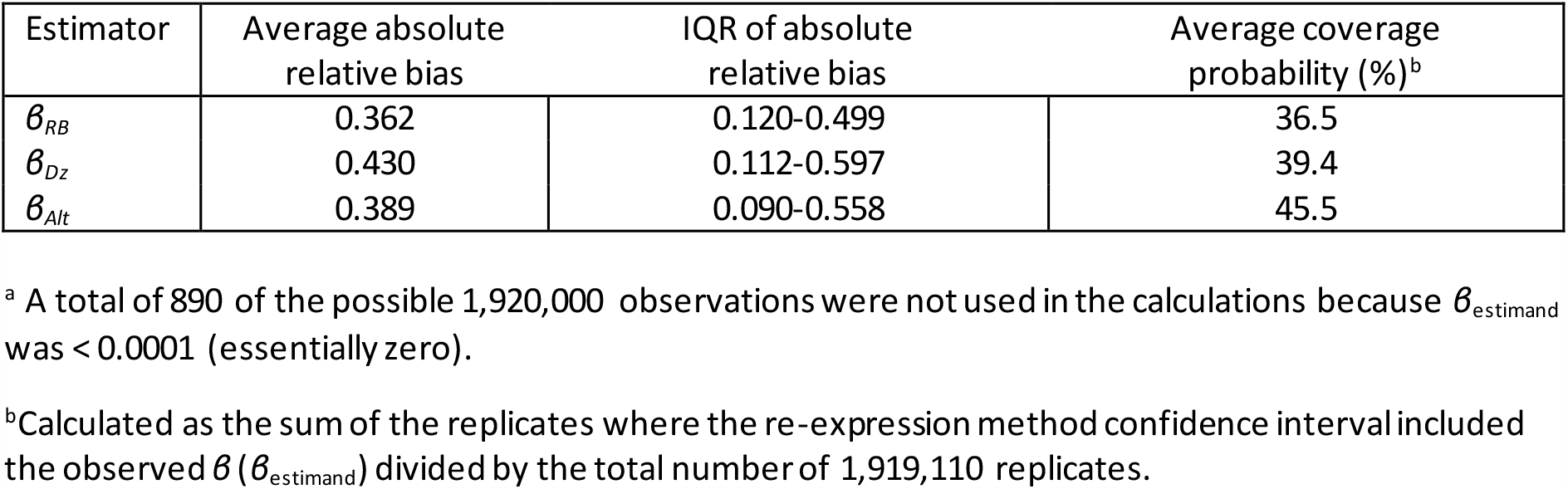
Performance measures based on simulations with all possible values for each parameter for a total of 960 simulation scenarios with 2000 simulations per scenario (n = 1,919,110).^a^

### 3.2 The real data and application of the estimators to it

We identified nine published analyses of data for which the raw data were publicly available and that met our criteria for selection (Table S3) (Abraham et al., 2020; Bulka et al., 2021; Cheang et al., 2021; Lee et al., 2020; Odebeatu et al., 2019; Pilkerton et al., 2018; Stein et al., 2016; Xu et al., 2020).

The results of our re-analyses were generally the same order of magnitude as those originally published (Table S3). The specific finding that we used in the analysis and its location in the original publication are listed in Supplementary Material Table S4, as are the median, quartiles, and mean of the exposure distributions, which were estimated in some cases as indicated by table footnotes. We identified six published analyses of data where the original authors presented regression results using exposure with and without a log-transformation (Table S5) (Apelberg et al., 2007; Chen et al., 2012; Darrow et al., 2013; Hamm et al., 2010; Steenland et al., 2018; Washino et al., 2009). Five of these were included in the assessment of validity by Dzierlenga et al. (2020) (Dzierlenga et al., 2020). Among the fifteen example studies, a variety of outcomes and exposure variables were examined, though in two thirds of the studies the exposure was a perfluoroalkyl substance (either PFOA or perfluorooctane sulfonic acid).

When *β* was re-expressed as if it had been fit to untransformed exposure data, the range in relative bias across all three estimators was -0.49 to 16.82 (Table 3) and the interquartile ranges in relative bias were relatively wide. In the comparison of results for specific studies across re-expression methods, the relative bias was, for most of the studies, similar across methods (Table 3). These were studies where the median exposure was > 4 units (Table S4) – as would be expected based on Figure 2D. For the Lee et al. (2020) and the two Xu et al. (2020) results (Lee et al., 2020; Xu et al., 2020), however, *β*_RB_ performed better than the other two methods. In these three instances, the median of the exposure variable was less than one, which was not the case for the other studies (see Supplementary Materials Table S4) and the *σ* was > 0.8 – which is the setting where the relative bias in *β*_RB_ was expected to be relatively small compared with the other estimators.

**Table 3:** Comparison of fitted and re-expressed *β* coefficients and relative bias in *β* for three methods of re-expression

| First author, year | $\beta_{\text{Estimand}}$ from analysis of raw data | $\beta_{\text{RB}}^a$ | Relative Bias<sup>b</sup> | $\beta_{\text{Dz}}$ | Relative Bias<sup>b</sup> | $\beta_{\text{Alt}}^c$ | Relative Bias<sup>b</sup> |
| --- | --- | --- | --- | --- | --- | --- | --- |
| Abraham, 2020 | -0.0636 log<sub>e</sub>/ ng·ml<sup>-1</sup> | -0.0324 | -0.49 | -0.0368 | -0.42 | -0.0384 | -0.40 |
| Apelberg, 2007 | -12.9 g/(ng/ml) | -11.7 | -0.10 | -13.2 | 0.03 | -13.2 | 0.03 |
| Bulka, 2021 | 0.015 log<sub>e</sub>(OR)/ ng·ml<sup>-1</sup> | 0.043 | 1.87 | 0.0512 | 2.41 | 0.0531 | 2.54 |
| Cheang, 2021 | 0.192 mg·dl/pmol·g<sup>-1</sup> | 0.335 | 0.74 | 0.352 | 0.83 | 0.353 | 0.84 |
| Chen, 2012 | -11.3 g/(ng/ml) | -16.3 | 0.44 | -17.9 | 0.58 | -17.8 | 0.58 |
| Darrow, 2013 | -2.3 g/(ng/ml) | -1.95 | -0.15 | -2.02 | -0.12 | -2.00 | -0.13 |
| Hamm, 2010 | 1.5 g/(ng/ml) | 3.66 | 1.44 | 3.92 | 1.62 | 3.85 | 1.57 |
| Lee, 2020 | 1.444 log<sub>e</sub>(OR)/μg·dl<sup>-1</sup> | 1.31 | -0.09 | 3.30 | 1.28 | 3.46 | 1.4 |
| Odebeatu, 2019 | 6.86x10<sup>-4</sup> log<sub>e</sub>(OR)/ng·ml<sup>-1</sup> | 0.0114 | 15.69 | 0.0122 | 16.82 | 0.0116 | 15.86 |
| Pilkerton, 2018 | -0.0049 log<sub>e</sub>/ng·ml<sup>-1</sup> | -0.0207 | 3.22 | -0.0302 | 5.17 | -0.0306 | 5.24 |
| Steenland, 2009 | 0.00105 log<sub>e</sub>/ ng·ml<sup>-1</sup> | 0.00116 | 0.11 | 0.00127 | 0.21 | 0.00126 | 0.20 |
| Stein, 2016 | -0.0039 log<sub>e</sub>/ng·ml<sup>-1</sup> | -0.00676 | 0.73 | -0.0069 | 0.77 | -0.00694 | 0.78 |
| Washino, 2009 | -10.94 g/(ng/ml) | -11.4 | 0.04 | -12.1 | 0.10 | -10.0 | -0.08 |
| Xu, 2020a | 0.513 log<sub>e</sub>(OR)/ ng·ml<sup>-1</sup> | 0.508 | -0.01 | 0.962 | 0.87 | 1.01 | 0.96 |
| Xu, 2020b | 29.3 mg·dl/ng·ml<sup>-1</sup> | 25.4 | -0.13 | 48.1 | 0.64 | 50.4 | 0.72 |
| Median |  |  | 0.11 |  | 0.77 |  | 0.78 |
| 25<sup>th</sup> percentile |  |  | -0.095 |  | 0.155 |  | 0.115 |
| 75<sup>th</sup> percentile |  |  | 1.09 |  | 1.45 |  | 1.485 |
| Minimum |  |  | -0.49 |  | -0.42 |  | -0.40 |
| Maximum |  |  | 15.69 |  | 16.82 |  | 15.86 |

<sup>a</sup> Using the notation of Rodriguez-Barranco et al., k = base of log transformation used; c = 1.
<sup>b</sup> Proportional difference between $\beta$ in column to the left compared with the one from the analysis of raw data, calculated using the same method as in previous table ((beta in column to left – beta from analysis of raw data)/beta from analysis of raw data). Note that the $\beta$ in column to the left was calculated using $\beta$ with different units in denominator than for the raw data analysis shown in the table.
<sup>c</sup> Let the log unit increment $l$ in untransformed units = $b^{(\log_b(\text{median}) + 0.5)} - b^{(\log_b(\text{median}) - 0.5)}$ , where $b = 2, e, \text{ or } 10$ , depending on the base. For observed $\beta_o$ with units of $\Delta y / \Delta \log_b(x)$ , to get re-expressed $\beta_r$ with units $\Delta y / \Delta x$ , calculate $\beta_r = \beta_o / l$ . If the units of $\beta_o$ are $\Delta y / \Delta x$ , to get $\beta_r$ with units $\Delta y / \Delta \log_b(x)$ , calculate $\beta_r = \beta_o \cdot l$ .

Our results for Odebeatu et al. (2019) and Pilkerton et al. (2018) were the ones with the greatest discrepancy between the re-expressed *β* coefficients and the *β* fitted to the untransformed exposure (Odebeatu et al., 2019; Pilkerton et al., 2018)(Table 3). This discrepancy suggested that there may have been observations that were influential, and that the influence was affected by whether the exposure had been log-transformed. Thus, we conducted an analysis of whether exclusion of influential points affected the accuracy of the re-expression. For comparison, similar analyses were conducted using the data from Cheang et al. (2021) and Xu et al. (2020) (dichotomous outcome), which showed smaller relative differences between the re-expressed and fitted *β*s. The analyses with and without the inclusion of especially influential points in the real data sets showed that the accuracy of the re-expression estimators was affected by their exclusion (Table S6). The relative bias was affected by the influential points more so for Odebeatu et al. (2019) and Pilkerton et al., (2019) than for Cheang et al. (2021) and Xu et al. (2020), but even with the exclusion of influential observations the re-expression methods still performed poorly.

When we used the regression equations (Table S3) to predict the relative bias in the estimators when applied to each of the real data examples, and then adjusted the re-expressed *β* to remove the bias, the adjusted *β*s, on average, showed less bias, but as before, the interquartile range of the adjusted relative bias was wide (Table 4).

**Table 4:** Comparison of fitted and re-expressed β coefficients and proportional difference in β for three methods of re-expression, adjusted by their OLS describing the relationship between sigma and relative bias.

| First author, year | $\beta_{\text{Estimand}}$ from analysis of raw data | $\beta_{\text{RB}}^a$ | Relative Bias<sup>b</sup> | $\beta_{\text{Dz}}$ | Relative Bias<sup>b</sup> | $\beta_{\text{Alt}}^c$ | Relative Bias<sup>b</sup> |
| --- | --- | --- | --- | --- | --- | --- | --- |
| Abraham, 2020 | -0.0636 log<sub>e</sub>/ng·ml<sup>-1</sup> | -0.0202 | -0.68 | -0.0214 | -0.66 | -0.0214 | -0.66 |
| Apelberg, 2007 | -12.9 g/(ng/ml) | -10.6 | -0.18 | -9.41 | -0.27 | -9.37 | -0.27 |
| Bulka, 2021 | 0.015 log<sub>e</sub>(OR)/ng·ml<sup>-1</sup> | 0.0339 | 1.26 | 0.0303 | 1.02 | 0.0301 | 1.01 |
| Cheang, 2021 | 0.192 mg·dl/pmol·g<sup>-1</sup> | 0.224 | 0.17 | 0.288 | 0.50 | 0.290 | 0.51 |
| Chen, 2012 | -11.3 g/(ng/ml) | -14.8 | 0.31 | -12.8 | 0.14 | -12.8 | 0.13 |
| Darrow, 2013 | -2.3 g/(ng/ml) | -1.69 | -0.27 | -1.56 | -0.32 | -1.56 | -0.32 |
| Hamm, 2010 | 1.5 g/(ng/ml) | 3.79 | 1.52 | 3.24 | 1.16 | 3.24 | 1.16 |
| Lee, 2020 | 1.444 log<sub>e</sub>(OR)/μg·dl<sup>-1</sup> | 0.957 | -0.34 | 1.79 | 0.24 | 1.79 | 0.24 |
| Odebeatu, 2019 | 6.86x10<sup>-4</sup> log<sub>e</sub>(OR)/ng·ml<sup>-1</sup> | 0.00374 | 4.45 | 0.00361 | 4.26 | 0.00364 | 4.30 |
| Pilkerton, 2018 | -0.0049 log<sub>e</sub>/ng·ml<sup>-1</sup> | -0.021 | 3.26 | -0.0232 | 3.74 | -0.0231 | 3.71 |
| Steenland, 2009 | 0.00105 log<sub>e</sub>/ng·ml<sup>-1</sup> | 0.00088 | -0.16 | 0.00090 | -0.14 | 0.00090 | -0.14 |
| Stein, 2016 | -0.0039 log<sub>e</sub>/ng·ml<sup>-1</sup> | -0.00527 | 0.35 | -0.00542 | 0.39 | -0.00542 | 0.39 |
| Washino, 2009 | -10.94 g/(ng/ml) | -11.6 | 0.06 | -9.34 | -0.15 | -9.40 | -0.14 |
| Xu, 2020a | 0.513 log<sub>e</sub>(OR)/ng·ml<sup>-1</sup> | 0.380 | -0.26 | 0.535 | 0.04 | 0.534 | 0.04 |
| Xu, 2020b | 29.3 mg·dl/ng·ml<sup>-1</sup> | 19.0 | -0.35 | 26.7 | -0.09 | 26.7 | -0.09 |
| Median |  |  | 0.06 |  | 0.14 |  | 0.13 |
| 25<sup>th</sup> percentile |  |  | -0.265 |  | -0.145 |  | -0.140 |
| 75<sup>th</sup> percentile |  |  | 0.805 |  | 0.760 |  | 0.760 |
| Minimum |  |  | -0.68 |  | -0.66 |  | -0.66 |
| Maximum |  |  | 4.45 |  | 4.26 |  | 4.30 |

<sup>a</sup> Using the notation of Rodriguez-Barranco et al., k = base of log transformation used; c = 1.
<sup>b</sup> Proportional difference between $\beta$ in column to the left compared with the one from the analysis of raw data, calculated using the same method as in previous table ((beta in column to left – beta from analysis of raw data)/beta from analysis of raw data). Note that the $\beta$ in column to the left was calculated using $\beta$ with different units in denominator than for the raw data analysis shown in the table.
<sup>c</sup> Let the log unit increment $l$ in untransformed units = $b^{(\log_b(\text{median}) + 0.5)} - b^{(\log_b(\text{median}) - 0.5)}$ , where $b = 2, e, \text{ or } 10$ , depending on the base. For observed $\beta_o$ with units of $\Delta y / \Delta \log_b(x)$ , to get re-expressed $\beta_r$ with units $\Delta y / \Delta x$ , calculate $\beta_r = \beta_o / l$ . If the units of $\beta_o$ are $\Delta y / \Delta x$ , to get $\beta_r$ with units $\Delta y / \Delta \log_b(x)$ , calculate $\beta_r = \beta_o \cdot l$ .

## 4. Discussion

In the simulations, the bias in each of the three estimators was evaluated in relation to the median of the exposure variable, the skewness in the exposure variable, the log base used to transform the exposure variable, the *β* in the modelgenerating the data, and the number of observations simulated. For all three re-expression methods, the relative bias was more positive as the skewness of the exposure distribution increased. The relative bias in *β*_RB_ was also determined by the median of the exposure distribution, and the relative bias in *β*_Alt_ was also affected by the base of the log used to transform the exposure variable. Although a few specific circumstances were found where the relative bias in a given re-expression method was lower, in general, when the skewness of *x* was large enough that a log transformation might be applied, the methods gave results that were sufficiently biased that their use would not be advisable. The results from applying the re-expression methods to real datasets generally agreed with those from the simulation, but their performance was worse than predicted based on the simulations. The poor performance in the real data was not much affected by the exclusion of influential observations. The especially poor performance of the re-expression methods in the case of the Odebeatu et al. (2019) data may have been due to the small size of the slope being re-expressed.

Rodriguez-Barranco et al. (2017) recognized the importance of skewness in causing bias in their estimator, though the degree of skewness in their simulations was not specified and only one median value was used. For the re-expression method proposed by Steenland et al. (2018), apparently it was assumed that if an exposure distribution had an upper bound near 10 units, their empirical re - expression method would be sufficiently accurate (Steenland et al., 2018). Our results suggested that the range of exposure was predictive of the validity of the re-expression only for the RB estimator. Using a re-expression method similar to Steenland et al.’s (Dzierlenga et al., 2020), Dzierlenga et al. (2020) conducted a validation study, using only five studies. Three of the studies in that validation had the least bias in the present study; their relatively small number of validation studies may have led to an overly optimistic appraisal of the method.

In this report we focused on re-expressing regression coefficients from linear models fit to a log-transformed exposure variable. We could have also addressed the opposite: re -expression of regression coefficients from linear models fit to the untransformed exposure variable. To simplify the manuscript, we did not address this opposite type of re-expression. In risk assessment, results based on untransformed exposure are of greatest use, hence our focus on expressing all results in arithmetic units.

The results of this assessment of validity have implications for systematic reviewers and meta-analysts considering or using these re-expression methods. The bias due to re-expression with the three methods evaluated was affected by the skewness of the exposure variable, and, for some estimators, the median exposure or the type of transformation used. Meta-analyses stratified by whether the key independent variable was log-transformed and synthesis of the stratified results without meta-analysis (McKenzie & Brennan, 2022) may be preferable to use of the re-expression methods studied here.

## Abbreviations

*β*_Estimand_: *β* coefficient from fitting *y* = *β*_Estimand_·*x* + *e* with an ordinary least squares (OLS) model
*β*_RB_: Estimator evaluated using the Rodriguez-Barranco method
*β*_Dz_: Estimator evaluated using the Dzierlenga method
*β*_Alt_: Estimator evaluated using the Alternative method
ADEMP: Aims, Data generating mechanism, Estimand, Methods, and Performance measures (Simulation study methodology)
DGM: Data generating mechanism
PFOA: Perfluorooctanoic acid

## Data Availability Statement

The data used in this manuscript are freely available from their respective publications or upon request from the original authors. R scripts/functions and data files for applying each of these three reexpression methods are available in the supplemental materials (Supplemental_Code.zip).

## Ethics Approval Statement

Not applicable

## Patient Consent Statement

Not applicable

## Permission to Reproduce Material from Other Sources

Not applicable

## Clinical Trial Registration

Not applicable

## Author roles

Dr. Linakis ran the re-expression algorithm on the examples and simulation datasets, conducted the analysis of factors that determine validity, computed results using the other re-expression methods, and contributed to the draft of the manuscript.

Ms. Van Landingham replicated the analysis of the example NHANES studies, conducted the influence analyses, and contributed to the draft of the manuscript.

Dr. Gasparini helped to critically evaluate and provide feedback on the methods and presentation of the results, and contributed to the draft of the manuscript.

Dr. Longnecker conceived of the study, designed it, obtained funding for it, and drafted the manuscript.

## Acknowledgments

Dr. Michael W. Dzierlenga provided critical comments on an early draft of the manuscript.

