## Supplemental Tables and Figures for "Re-expressing coefficients from regression models for inclusion in a meta-analysis"

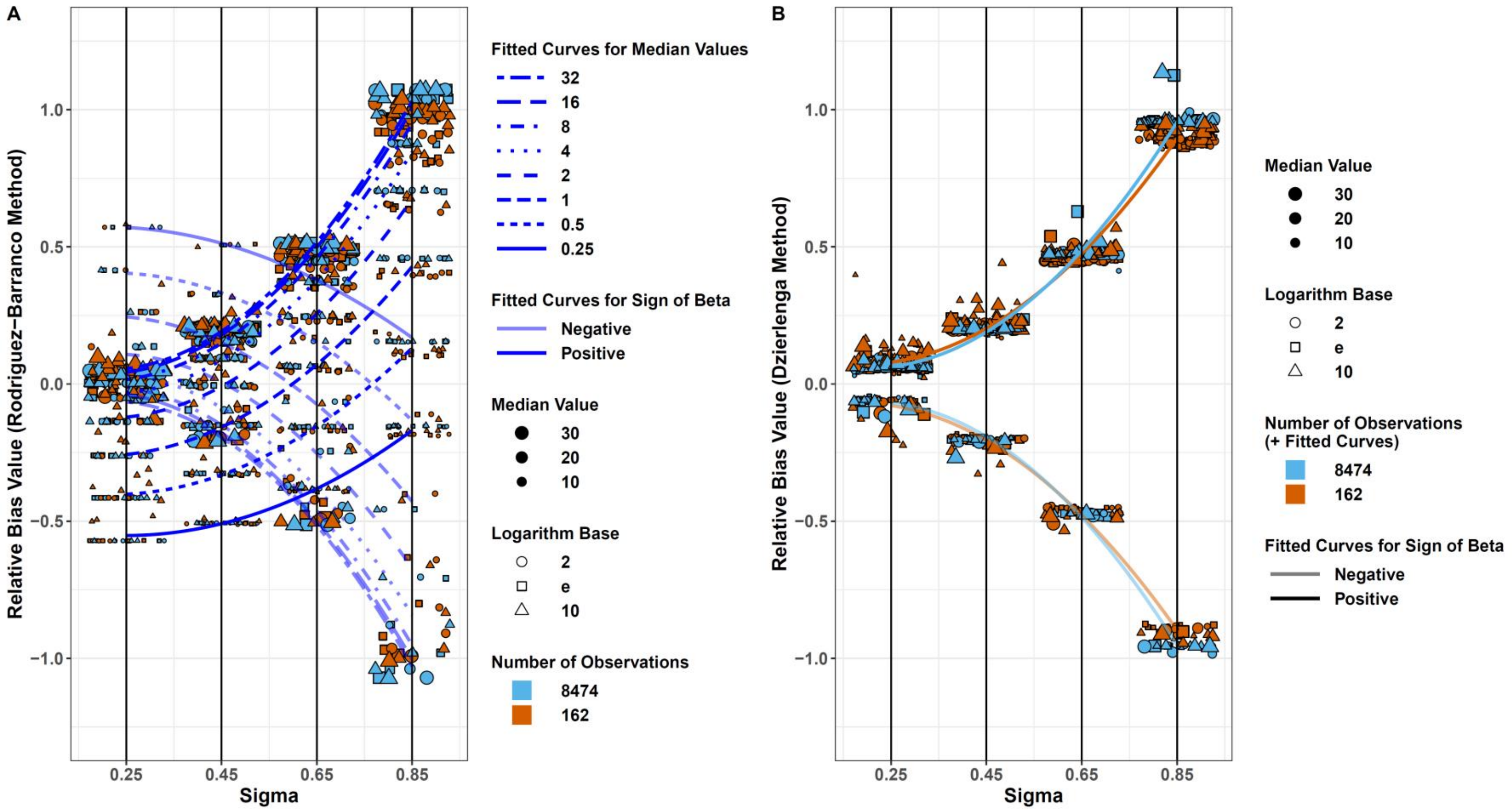
A
Relative Bias Value (Rodriguez−Barranco Method)
1.0
0.5
0.0
−0.5
−1.0
0.25
0.45
0.65
0.85
Sigma
Fitted Curves for Median Values
32
16
8
4
2
1
0.5
0.25
Fitted Curves for Sign of Beta
Negative
Positive
Median Value
30
20
10
Logarithm Base
2
e
10
Number of Observations
8474
162
B
Relative Bias Value (Dzierlenga Method)
1.0
0.5
0.0
−0.5
−1.0
0.25
0.45
0.65
0.85
Sigma
Median Value
30
20
10
Logarithm Base
2
e
10
Number of Observations
(+ Fitted Curves)
8474
162
Fitted Curves for Sign of Beta
Negative
Positive

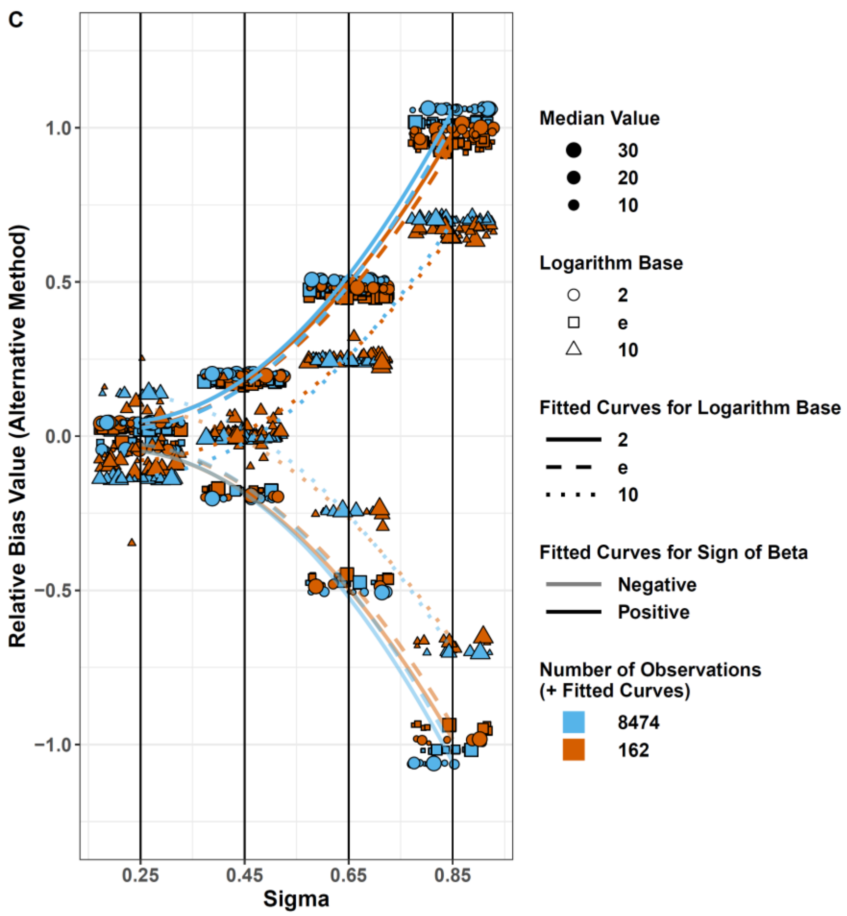

Figure S1: Plots of relative bias as a function of skewness (σ) in the exposure x, by type of estimator, including the scenario where $\beta_{DGM} < 0$. Individual points represent the average result ($n_{sim} = 2000$) for each simulation scenario. A total of 890 of the possible 1,920,000 observations (960 scenarios x 2000 simulations) were not used in the calculation of the average results because $\theta_{estimand}$ was < 0.0001 (essentially zero). Lines represent quadratic fits to the data for a specified prediction equation and set of values of independent variables (see text). Note that data have been artificially spread along the x-axis for visualization purposes, all actual x-values are the closest black vertical line (0.25, 0.45, 0.65, or 0.85). Figures A-C show points for 960 simulations. A) Rodriguez-Barranco estimator, B) Dzierlenga estimator, and C) Alternative estimator.

Table S1: Coefficients from ordinary least squares models of relative bias, by re-expression method. Each method was described by a quadratic fit ($ax^2+bx+c$) with additional predictor variables as described in the table.

| Method | Parameter | Coefficient | Standard Error | *p*-value | RMSE | Adjusted $R^2$ |
|---|---|---|---|---|---|---|
| **Rodriguez-Barranco Method ($\beta_{RB}$)** | Intercept (c) | -0.13 | 0.0596 | 0.026 | 0.230 | 0.706 |
| | Linear sigma term (b) | -0.95 | 0.2342 | <0.001 | | |
| | Quadratic sigma term (a) | 1.87 | 0.2085 | <0.001 | | |
| | Median | 0.01 | 0.0021 | 0.001 | | |
| | Interaction term between median and sigma (μ:σ) | 0.02 | 0.0036 | <0.001 | | |
| **Dzierlenga Method ($\beta_{Dz}$)** | Intercept (c) | 0.18 | 0.0076 | <0.001 | 0.028 | 0.993 |
| | Linear σ term (b) | -0.84 | 0.0283 | <0.001 | | |
| | Quadratic σ term (a) | 2.00 | 0.0251 | <0.001 | | |
| | Number of observations in simulated study (nobs) | -7.02E-06 | 6.40E-07 | <0.001 | | |
| | logbase (logbase = 10) | 0.0167 | 0.0025 | <0.001 | | |
| | logbase (logbase = 2) | -0.0026 | 0.0025 | 0.29 | | |
| | $\beta_{DGM}$ | 0.0002 | 8.39E-05 | 0.004 | | |
| | Median | 0.0005 | 9.68E-05 | <0.001 | | |
| | Interaction term between nobs and sigma (nobs:σ) | 1.51E-05 | 1.08E-06 | <0.001 | | |
| **Alternative Method ($\beta_{Alt}$)** | Intercept (c) | 0.13 | 0.0071 | <0.001 | 0.024 | 0.996 |
| | Linear σ term (b) | -0.90 | 0.0247 | <0.001 | | |

| | | | | |
|---|---|---|---|---|
| | Quadratic σ term (a) | 2.20 | 0.0214 | <0.001 |
| | logbase (logbase = 10) | -0.05 | 0.0056 | <0.001 |
| | logbase (logbase = 2) | 9.41E-03 | 0.0056 | 0.091 |
| | number of observations in simulated study (nobs) | -7.21E-06 | 5.46E-07 | <0.001 |
| | Interaction term between σ and logbase = 10 (logbase10: σ) | -0.27 | 0.0094 | <0.001 |
| | Interaction term between σ and logbase = 2 (logbase2: σ) | 0.04 | 0.0094 | <0.001 |
| | Interaction term between σ and nobs (σ:nobs) | 1.56E-05 | 9.20E-07 | <0.001 |

Confidential

Table S2: Comparison of model fits ($R^2$) when using each simulation scenario with $\beta_{DGM} > 0$ (n = 768) or using each observation with $\beta_{DGM} > 0$ (n = 768 * 2000)

| **$r^2$ for:** | **$\beta_{RB}$** | **$\beta_{Dz}$** | **$\beta_{Alt}$** |
|---|---|---|---|
| **n = 768** | 0.706 | 0.992 | 0.996 |
| **n = 768 * 2000** | 0.709 | 0.992 | 0.996 |

Table S3. Published analyses of an outcome in relation to a biomarker-based measure of environmental exposure, with raw data available

| 1st author, year | Outcome | Type of outcome[a] | Exposure[b] | Original unit of exposure | Result presented by original authors[c] | Our result[c] (re-analysis of raw data) |
|---|---|---|---|---|---|---|
| Bulka, 2021 | Herpes Simplex Virus 2 | D | PFOA | $Log_2$ | 1.11 (1.05, 1.17) | 1.11 (1.05, 1.17) |
| Lee, 2020 | Infertility | D | Cadmium | $Log_2$ | 1.8 (1.1, 3.1) | 1.8 (1.1, 3.1) |
| Odebeatu, 2019 | Asthma | D | Mono-benzyl phthalate (urine) | $Log_{10}$ | 1.50 (1.09, 2.08) | 1.50 (1.08, 2.08) |
| Xu, 2020 | CVD[d] | D | Isopentanaldehyde | ng/ml | P<0.001 | 1.67 (1.17, 2.25) |
| Xu, 2020 | Triglycerides (mg/dl) | C | Isopentanaldehyde | ng/ml | 25.0 (4.8, 45.1) | 29.3 (14.1, 44.6) |
| Stein, 2016 | Mumps IgG[e] (%Δ) | C | PFOS | $Log_2$ | -7.4 (-12.8, -1.7) | -10.3 (-19.3, -0.024) |
| Pilkerton, 2018 | Rubella IgG[e] (%Δ) | C | PFOA | $Log_2$[f] | -8.9 (-16.9, -0.2)[f] | N.A. |
| Cheang, 2021 | Triglycerides (mg/dl) | C | Glycidamide[g] | $Log_2$ | 11.4 (5.1, 17.7) | 9.65 (1.62, 17.7) |
| Abraham, 2020 | Ln(Hib[h] IgG[e]) | C | PFOA | $Log_2$ | -0.3887[i] (-0.6957, -0.0817) | N.A. |

[a] C = continuous, D = dichotomous

[b] Measured in serum unless noted otherwise

[c] Results shown are from regression analyses. For dichotomous outcomes, these are odds ratios (and 95% confidence intervals). For continuous outcome, the results are either regression coefficients, or regression results re-expressed as percent difference (%Δ) in outcome per unit exposure

[d] CVD, cardiovascular disease

[e] IgG, immunoglobulin G

[f] The original units were quartiles; we re-analyzed the data to get the original result shown, in percent change in Rubella antibody per $log_2$ increase in PFOA. See Crawford et al. (reference), Supplementary Material, Section X for an account of the re-analysis.

[g] As reflected by concentration of hemoglobin adduct of glycidamide (HbGA)

[h] Hib, Hemophilus Influenza

[i] Our analysis of the Abraham data. See Crawford et al. (submitted, 2021), Supplementary Material, Section S7 for an account of the re-analysis.

Table S4. Additional details about the 15 example studies

| Study, Year | Specific Finding (Location, Outcome) | Exposure Distribution | | | | |
|---|---|---|---|---|---|---|
| | | Median | 1st Quartile | 3rd Quartile | σ (lognormal distribution) | Mean |
| Abraham 2020 | Our analysis, Hib IgG | 14.3[a] | 6.70 | 19.3 | 0.78 | 16.8 |
| Apelberg, 2007 | Table 3 (Fully Adjusted), Birth Weight (g) | 5.00[b] | 3.40 | 7.90 | 0.62 | 5.43[f] |
| Bulka, 2021 | Table 3 (20-49 y), HSV 2 | 2.77[a] | 1.67 | 4.6 | 0.75 | 3.0[f] |
| Cheang, 2021 | Table 3, Triglycerides (mg/dL) | 38.7[a] | 29.4 | 55.2 | 0.47 | 41.1[f] |
| Chen, 2012 | Table 3 (Adjusted), Birth Weight (g) | 5.94[e] | 3.94 | 8.94 | 0.61 | 6.27[f] |
| Darrow, 2013 | Table 6 (Adjusted All Births, Per in unit increase), Birth Weight (g) | 13.9[b] | 9.5 | 19.7 | 0.54 | 14.4[f] |
| Hamm, 2010 | Table 5 (Hamm, PFOS), Birth Weight (g) | 7.80[d] | 5.70 | 10.7 | 0.47 | 8.07[f] |
| Lee, 2020 | Table 2 (Model 2), Infertility | 0.240[a] | 0.14 | 0.43 | 0.83 | 0.270[f] |
| Odebeatu, 2019 | Figure 1a (MBzP), Asthma | 12.3[b] | 5.00 | 27.3 | 1.26 | 14.9[f] |
| Pilkerton, 2018 | Table 4, Rubella (%Δ) | 4.30[a] | 3.00 | 6.3 | 0.55 | 6.00 |
| Steenland 2009 | Table 4, Total Cholesterol | 20.2[c] | 13.6 | 29.3 | 0.57 | 22.4 |
| Stein, 2016c | Table 2, Mumps (%Δ) | 22.2[a] | 15.35 | 30.8 | 0.52 | 22.8[f] |
| Xu, 2020 | Table 2 (Model 1), CVD | 0.521[a] | 0.346 | 1.03 | 0.81 | 0.632[f] |
| Xu, 2020 | Table 4 (Model 2), Triglycerides (mg/dL) | 0.521[a] | 0.346 | 1.03 | 0.81 | 0.632[f] |
| Washino, 2009 | Table 5 (Fully Adjusted), Birth Weight (g) | 5.20[b] | 3.40 | 7.00 | 0.53 | 5.20[f] |

[a]Median and IQR calculated from raw data

[b]Median and IQR pulled from publication

[c]Median from publication and IQR adjusted from Steenland 2010 by subtracting 0.6 (difference between medians in studies)

[d]Median from publication and IQR adjusted from Lind 2017, by subtracting 0.3 (difference between medians in studies)

[e]Median from publication and IQR adjusted from Chen 2017 by adding 0.24 (difference between medians in studies)

[f]Mean estimated from median and IQR using the formula from Wan 2014: (Q1+Median+Q3)/3

Table S5. Published analyses of an outcome in relation to a biomarker-based measure of environmental exposure, where the original authors presented results using exposure with and without a log-transformation

| 1st author, year | Outcome | Type of outcome | Exposure | Logarithmic (β in units/log(ng/ml)) Result reported by original authors[a] | Untransformed (β in units/(ng/ml)) Result reported by original authors[b] |
|---|---|---|---|---|---|
| Apelberg, 2007 | Birth weight (g) | C | PFOS | -69 (-149, 10) | -12.9 (-27.8, 2) |
| Chen, 2012 | Birth weight (g) | C | PFOS | -110.2 (-176.0, -44.5) | -11.30 (-17.40, -5.20) |
| Darrow, 2013 | Birth weight (g) | C | PFOS | -29 (-66, 7) | -2.3 (-4.8, 0.3) |
| Hamm, 2010 | Birth weight (g) | C | PFOS | 31.3 (-43.3, 105.9) | 1.5 (-7.6, 10.6) |
| Steenland, 2009 | Ln(Serum cholesterol (mg/dl)) | C | PFOS | 0.0266 (0.0239, 0.0293) | 0.00105 (0.0009, 0.0012) |
| Washino, 2009 | Birth weight (g) | C | PFOS | -148.8 (-297.0, -0.5) | -10.94 (-22.9, 1.10) |

[a] Values shown are for $\log_e$ transformations, except for Washino et al, for which a $\log_{10}$ transformation was used. Values in parentheses are 95% confidence intervals.
[b] Values for Washino et al. (2009) and Chen et al. (2012) in this column were presented in Verner et al. (2015).

Table S6

| 1st author, year | Proportional difference with influential observations | Proportional difference with no influential observations |
|---|---|---|
| Odebeatu 2019 | 16.82 | 19.58 |
| Pilkerton 2018 | 5.17 | 2.18 |
| Cheang 2021 | 0.84 | 1.11 |
| Xu 2020a | 0.88 | 1.38 |
